# Mapping the circulating proteome across neurodegeneration: A harmonized, consortium-scale framework for uncovering molecular pathophysiology

**DOI:** 10.64898/2025.12.09.693329

**Authors:** Caitlin A. Finney, Lijun An, Laura M. Winchester, Jacob Vogel, Heather M. Wilkins, Jeffrey M. Burns, Russell H. Swerdlow, Chad Slawson, Jeffrey D. Rothstein, Global Neurodegeneration Proteomics Consortium (GNPC), Michael W. Lutz, Rowan Saloner, Artur Shvetcov

## Abstract

Large-scale plasma proteomics offers unprecedented opportunities to investigate the systemic biology of neurodegeneration, yet technical heterogeneity, site-specific artifacts, and clinical confounding remain major barriers to reproducible discovery. Leveraging data from 13,733 individuals with Alzheimer’s disease (AD), Parkinson’s disease (PD), frontotemporal dementia (FTD), Parkinson’s disease dementia (PDD), amyotrophic lateral sclerosis (ALS), and non-impaired controls in the Global Neurodegeneration Proteomics Consortium (GNPC), we present a scalable and generalizable analytical framework for harmonizing and interpreting consortium-scale proteomic datasets. Using a high-dimensional perturbation framework, we systematically benchmark five commonly used batch correction methods across a range of realistic confounding structures, including site-disease imbalance, nonlinear effects, and heteroskedasticity. Empirical Bayes modelling via limma consistently emerged as the most robust method, optimally balancing removal of site-related technical variance with retention of disease-relevant biological signal. On this harmonized foundation, we resolve neurodegenerative disease plasma signatures, including a shared immune-metabolic axis in AD and PD, neuromuscular disruption in ALS, and proteostatic imbalance in PD. Tissue and cell-type enrichment highlight widespread immune-endocrine involvement in AD and hematopoietic activation in PD. Demographically matched analyses nominate distinct, candidate biomarkers across diseases, including lipid, redox, and complement factors in AD, lysosomal and cytoskeletal proteins in PD, and muscle-derived markers in ALS. This study establishes a scalable analytical framework for integrating real-world proteomic data and provides a disease-resolved catalogue of circulating signatures to inform biomarker development and targeted intervention across neurodegenerative diseases.

## Introduction

Neurodegenerative diseases, including Alzheimer’s disease (AD), Parkinson’s disease (PD), frontotemporal dementia (FTD), and amyotrophic lateral sclerosis (ALS), represent an escalating global health crisis driven primarily by demographic aging ^1^. By 2050, the global population over 60 years is expected to surpass 2.1 billion, with more than 420 million individuals over 80 ^2^. Neurodegenerative conditions are now among the leading causes of morbidity and mortality worldwide, placing profound personal and socioeconomic burden on patients, caregivers, and healthcare systems ^1,3,4^. Although major advances have refined our understanding of disease mechanisms, effective disease-modifying therapies remain limited. A central barrier is the lack of accessible, reproducible biomarkers that can detect early disease, resolve biological heterogeneity, guide therapeutic stratification, and monitor progression. Plasma biomarkers, in particular, hold transformative potential for population-level screening, precision clinical trials, and scalable implementation across diverse healthcare settings.

To meet this need, large-scale consortia have emerged to aggregate multi-omic datasets and unify analytical approaches across independent research centers. These efforts enhance statistical power, reproducibility, and translational reach. Notable exemplars include the Clinical Proteomic Tumor Analysis Consortium (CPTAC), which established proteogenomic standards across cancer centers ^5,6^, and the Accelerating Medicines Partnership (AMP), spanning AD ^7^, PD ^8^, schizophrenia ^9^, ALS ^10^, and immune-mediated diseases ^11–15^. The UK Biobank Pharma Proteomics Project further demonstrated the power of population-scale plasma proteomics for linking genetic variation to proteomic regulation ^16^.

More recently, the Global Neurodegeneration Proteomics Consortium (GNPC) has generated a uniquely comprehensive multi-site plasma proteomic resource across AD, PD and Parkinson’s disease dementia (PDD), FTD, and ALS, spanning more than 18,000 participants across 23 international sites ^17^. A key feature of GNPC v1 is its use of the aptamer-based SomaScan platform, enabling high-throughput quantification of thousands of proteins across broad dynamic ranges, including low-abundance species rarely detected by mass spectrometry ^18–22^. This provides unprecedented opportunities for mechanistic discovery, biomarker development, and therapeutic target prioritization across neurodegenerative diseases.

Realizing the vast potential of multi-site proteomics requires analytic strategies capable of rigorously harmonizing data across cohorts with samples collected under different protocols and run on separate assay batches. Real-world, multi-site plasma datasets such as GNPC are characterized by substantial technical and demographic heterogeneity ^23–26^. Batch effects arising from sample handling, platform calibration, or site-specific protocols can produce spurious associations, inflate classification performance, and compromise reproducibility ^24,26^. Yet overly aggressive correction may suppress biologically meaningful variance, particularly when technical and clinical factors are confounded, disproportionately affecting signals related to sex, ancestry, or disease subtype ^23,24^. Existing correction approaches, ranging from regression-based adjustment to empirical Bayes methods, perform inconsistently across confounded, real-world designs ^25–27^, underscoring the need for principled, data-driven benchmarking frameworks.

Here, we address this critical gap by systematically evaluating batch correction strategies within the GNPC dataset and establishing a reproducible framework for large-scale multi-site plasma proteomic integration. Building on the foundational GNPC resource ^17^, we benchmark correction approaches from site-level normalization to empirical Bayes harmonization, quantifying their effects on variance structure, biological signal retention, and recovery of diagnosis-associated protein changes. We further demonstrate how harmonized multi-site proteomic data can be leveraged to reveal conserved molecular signatures across neurodegenerative diseases, identify tractable therapeutic targets, and derive clinically actionable biomarker candidates. By integrating methodological rigor with mechanistic and translational discovery, this work provides both a generalizable analytical blueprint for consortium-scale proteomics and a high-confidence set of plasma biomarkers and pathways with relevance for early detection, patient stratification, and therapeutic development in neurodegeneration.

## Methods

### Global Neurodegeneration Proteomics Consortium (GNPC) cohort

The GNPC cohort is the world’s largest combined plasma proteomic dataset for individuals with different neurodegenerative diseases and non-impaired controls from clinical cohorts across the US, UK, and Europe ^17^. In the current study, we included 13,733 individuals from the GNPC cohort, with n = 8,372 non-impaired (NI) controls, 4,217 Alzheimer’s disease (AD), 175 frontotemporal dementia (FTD), 185 Parkinson’s disease dementia (PDD), 540 Parkinson’s disease (PD), and 244 amyotrophic lateral sclerosis (ALS) cases (Supplementary Table 1). For all individuals, we only used the first plasma measurement recorded from the contributing site and removed plasma measurements from subsequent visits. Individuals were diagnosed based on diagnostic criteria from each contributing cohort, as described elsewhere ^17^. Dementia thresholds for AD and PDD patients was further assessed using a clinical dementia rating (CDR) score of ≥ 1, Mini-Mental State Exam (MMSE) score of < 24, and/or Montreal Cognitive Assessment (MOCA) score of < 23 ^28,29^. Most individuals with AD had cognitive impairment scores indicative of mild or moderate AD (Supplementary Table 1). Non-impaired controls were identified locally by the contributing site as asymptomatic for neurodegenerative disease and no cognitive impairment. Individuals with a labelled diagnosis of mild cognitive impairment (MCI), or who did not meet CDR, MMSE, or MOCA thresholds for either AD or non-impaired control, were excluded from the present study, as MCI is not necessarily reflective of underlying neurodegenerative processes or disease. Participants from each of the included contributing sites in the GNPC cohort provided written informed consent and studies were approved by the relevant institution’s ethics committee ^17^.

### Plasma Proteomics

Individuals from across the contributing GNPC contributing sites provided plasma samples at a single timepoint. Plasma proteomics was done using SomaLogic’s SomaScan v4.1 assay that measures approximately 7,000 proteins. The SomaScan assay uses aptamer-based technology called slow off-rate modified aptamers (SOMAmers). These contain chemically modified nucleotides that bind with high specificity and affinity to target proteins ^18^. Final proteomic data provided by SomaLogic has been standardized, normalized, and calibrated via an Adaptive Normalization by Maximum Likelihood (ANML) pipeline, with protein measurements provided in relative fluorescent units (RFU). Prior to integration into the GNPC cohort dataset, aptamers were mapped to Uniprot as described elsewhere ^17^.

### Initial exploration of the plasma proteomic dataset

Principal component analysis (PCA) is one of the most frequently used tools in high-dimensional data exploration to assess technical variation in datasets. We therefore first used a PCA to understand the global structure of the data across all proteins and visualised clustering by diagnostic group and by contributing site. Individual samples were not excluded based on visual or distributional extremity unless clear evidence of technical artefact or data corruption was present (e.g. extreme missingness, biologically implausible values). No individual samples were excluded from analyses, as none were explicit outliers showing evidence of technical artefact or biologically implausible values. Observed dispersion was interpreted as plausible and consistent with expected heterogeneity in large multi-center cohorts. Standard statistical outlier detection methods were not applied, as they can erroneously flag valid samples in high-dimensional, multi-batch datasets ^30^. PCA plots were made using ‘ggplot2’ ^31^ in R (v4.4.1).

### Batch correction methods

We evaluated five common batch correction methods. These included per-site z-score standardization, linear model (LM) residual using site as a covariate, ComBat, ComBat(bal) which is a variant of ComBat excluding sites that are severely confounded by diagnosis, and limma. Biological and demographic covariates such as age and sex were deliberately omitted, because our goal at this stage was not to adjust for biological confounders, but to benchmark how effectively each correction method removes unwanted technical variation while preserving biologically meaningful outcome signal (e.g. AD vs. control). Including these covariates would have required specifying additional, overlapping sources of biological signal (e.g. age and sex effects and their interactions with site and outcome), making it more difficult to attribute changes in our performance metrics specifically to batch handling rather than to covariate adjustment. Z-score standardization is a simple within-batch scaling method where each protein is normalized to have zero mean and unit variance per site. In LM residual, each protein is fitted to a linear regression model with contributing site as a covariate, and residuals are extracted. This approach removes variance linearly attributable to site and assumes additive, linear effects. ComBat is a widely used empirical Bayes approach that adjusts for both additive and multiplicative batch effects. It models each protein’s expression as a function of batch and other covariates such as outcome and sex and applies empirical Bayes shrinkage to stabilize parameter estimates across proteins ^32^. We also developed a variation of ComBat, ComBat(bal), where sites confounded by diagnosis were excluded *a priori* to using ComBat. Specifically, we removed sites that only contributed one type of participant (e.g. only non-impaired controls or only neurodegenerative disease cases) for a total of five removed sites. Finally, limma was originally developed for microarray data and also uses a linear modelling framework with an empirical Bayes variance moderation.

Unlike ComBat, limma fits feature-wise linear models and then uses empirical Bayes moderation to stabilise protein level variance estimates. Variances for individual proteins are shrunk towards a common mean-variance trend estimated from all proteins, yielding more stable test statistics and higher power. ^33^.

For each method, we compared the global variance structure of corrected data with raw data. Reduction in the proportion of variance explained by leading PCs (e.g. PC1) was used as an initial indicator of site-related correction. To assess whether site separation persisted in multivariate space, we computed the median silhouette width over the top 10 PCs using site labels as cluster assignments. Silhouette values range from +1 (tight, well-separated clusters), through 0 (random mixing), to -1 (samples closer to other clusters than their own), providing an interpretable geometric measure of clustering strength.

### Random effects modelling of site- and outcome-related variance

To quantify residual technical and biological variance, we applied linear mixed-effects models with random intercepts for study site and diagnostic outcome (AD vs. control). This approach partitions total protein-level variance into components attributable to technical and biological sources, enabling estimation of site- and outcome-related effects while accounting for other sources of variability, and yielding per-protein variance decomposition profiles. For each correction method, we visualised the distributions of per-protein site and outcome variance and summarized the median variance attributable to each component. Retained outcome variance was quantified as the proportion of variance attributable to diagnostic outcome after correction, expressed relative to the corresponding estimate in the raw (uncorrected) dataset. Although this does not represent recovery of a known ground truth, it provides a relative measure of how well each method preserves biologically meaningful disease-related variance while mitigating technical artefacts.

### Perturbation study

To evaluate robustness across diverse analytical conditions, we conducted a large-scale perturbation study. We repeatedly sampled subsets of 2,000 participants and 1,000 proteins (100 iterations) from the full dataset, thereby preserving the complex covariance structure of the real data. Into these subsets we introduced controlled modifications to systematically vary several parameters. These included effect size of the injected outcome signal (0, 0.5, 1, 2), degree of batch-outcome confounding (none, moderate, and severe), presence of batch heteroskedasticity (True, False), nonlinearity of batch effects (True, False), and signal sparsity (0, 0.05, 0.1) to form 144 unique parameter scenarios. When heteroskedasticity is true, this introduces site-specific variance inflation to mimic heteroskedasticity across sites. For nonlinearity of batch effects, when true a site-specific nonlinear transformation is applied via a sine distortion to all protein values. Signal sparsity controls the proportion of proteins receiving the injected signal. Across each of the modification scenarios, we compared the five correction methods. Performance metrics included variance decomposition estimates for site and outcome using random effects modelling and a signal recovery rate defined as fraction of spiked proteins recovered among top-ranked hits. This framework enabled benchmarking of batch correction methods across a range of confounding structures and data-generating conditions, providing an integrated view of their relative strengths and limitations.

### Differential abundance analysis

Differentially expressed plasma proteins between individuals with a neurodegenerative disease and NI controls were identified using a dual-model framework combining pooled and meta-analytic approaches. This was coupled with comprehensive quality control metrics to assess cross-site replicability and flag potential artefacts arising from site-specific biases. We first applied a limma-based linear model to the batch-corrected expression matrix. For each protein, we computed moderated t-statistics, log_2_ fold changes (logFC), and associated *p*-values for the disease versus control comparison, with significance assessed using false discovery rate (FDR) correction. To further assess the robustness of differentially expressed proteins across collection sites, we performed site-specific differential expression analyses for all sites with at least 10 neurodegenerative disease and 10 control samples. Independent models were fitted for each site to estimate logFC and standard errors, which were then combined using inverse-variance weighted fixed-effects meta-analysis. Significance was defined both at nominal (*p* < 0.05) and FDR-adjusted thresholds.

A series of quality control metrics were computed for each protein to distinguish biologically meaningful signals from artefacts. First, we calculated the maximum site weight, the proportion of total inverse-variance weight contributed by the most influential site in the meta-analysis. Proteins with maximum site weight exceeding 0.6 were flagged as site-dominant and considered potentially sensitive to cohort-specific effects. Second, we assessed directional consistency by calculating the opposing weight proportion, defined as the proportion of total weight contributed by sites with effect directions opposite to the overall meta-analytic estimate. Proteins with opposing weight proportion ≥ 0.3 were flagged as directionally discordant, indicating inter-site heterogeneity. Third, we performed leave-one-site-out (LOSO) sensitivity analyses for all proteins with contributions from at least three sites. Proteins were flagged as fragile if omitting any single site reversed the direction of the meta-analytic effect or changed its significance from *p* < 0.05 to *p* > 0.05. Directional reversals were interpreted as evidence of site-driven instability, whereas loss of significance without reversal likely reflected marginal statistical power rather than genuine directional inconsistency. Based on pooled and meta-analytic significance, effect direction consistency, and the quality control metrics above, each protein was assigned to one of five groups (Table 1). Group 1 (high confidence) included proteins significant after correction with FDR < 0.05 in both pooled and meta-analyses, no signs of site dominance, directional discordance, or LOSO fragility, and consistent direction of effect. Group 2 (supported but underpowered) met the same criteria, but with nominal rather than FDR-corrected significance in the meta-analysis. Group 3 (likely site-driven) included proteins with high maximum site weight, substantial directional discordance, or LOSO fragility. Group 4 (ambiguous) included proteins significant after FDR significance correction in either model but with unresolved artefactual signals. Group 5 (background) comprised all proteins not meeting significance in either analysis.

**Table 1.** Summary of assigned protein labels in the current study.

| Label | Definition | Interpretation |
| --- | --- | --- |
| High confidence | Proteins passing both meta-analysis and pooled-analysis significance thresholds (adj. $p < 0.05$), showing consistent effect direction, robust to LOSO sensitivity analysis, and not dominated by a single site | Strong, reproducible signals likely to reflect disease biology |
| Supported but underpowered | Proteins significant in either meta-analysis or pooled analysis (but not both), with consistent direction and no signs of site dominance or instability | Potentially true effects limited by statistical power, particularly in smaller diagnostic groups |
| Likely site-driven | Proteins with statistical significance but failing robustness checks (e.g. strongly influenced by one site, or unstable under LOSO including flip in direction or loss of significance) | Caution warranted; effects may be driven by site-specific biases |
| Ambiguous | Proteins with conflicting results (e.g. direction flip between meta and pooled analysis, or direction disagreement across sites), or partially significant but unstable findings. | Inconclusive due to heterogeneity, borderline effects, or inconsistencies |
| Background | Proteins failing all significance thresholds (adj $p > 0.05$ in both meta and pooled analyses) and showing no evidence of meaningful or robust differential expression | No signal detected in current data |

### Sample size, age, and sex-matched analyses

To complement the primary analysis, we implemented a secondary, more conservative strategy focused on translational robustness. To control for demographic confounding across diagnostic groups, we conducted 1:1 case-control matching by age and sex for each disease group comparison against NI controls. Matching was performed using the ‘MatchIt’ package (v4.5.2) in R ^34^, specifying exact matching on sex and nearest-neighbour matching on age within a 2-year calliper. NI controls were drawn from a large pool of 8,372 samples, allowing stringent demographic alignment while maintaining balanced group sizes across most comparisons. By prioritizing demographic comparability over sample size, this approach minimizes residual confounding and strengthens the interpretability of disease-specific molecular signatures, particularly in smaller or heterogeneous cohorts.

### Pathway and tissue-specific enrichment analyses

Functional enrichment analyses of the identified plasma proteins were done using NetworkAnalyst (v3.0) ^35–37^. Here, generic protein-protein interactions were identified using first and zero order networks from the STRING interactome ^38^. A ‘high confidence’ score cutoff of 700 was used and experimental evidence was required for all protein-protein interactions identified. Network enrichment for biological processes and molecular functions was done using the PANTHER classification system ^39^ and Kyoto Encyclopedia of Genes and Genomes (KEGG) ^40^ was used for pathways. For each biological process, cellular component, and pathway, the false discovery rate (FDR) is reported. Protein-drug interaction enrichment analyses were also done in NetworkAnalyst (v3.0) ^35–37^ using information collected from the DrugBank database (v5.0) ^41–43^ and a minimum network. A linear bipartite graph was created in NetworkAnalyst and degree and betweenness for each pharmacologically active agent reported.

To contextualize the biological relevance of differentially expressed proteins identified in the batch effect correction analysis, we developed and applied a multi-scale enrichment pipeline implemented in the custom-built ‘xEnrich’ R package (https://github.com/Art83/xEnrich). This framework enables systematic investigation of gene or protein set enrichment across both bulk tissue level and single cell resolution datasets.

Briefly, the protein list was first subjected to enrichment analysis using immunohistochemistry microarray-based expression data from the Human Protein Atlas (HPA) ^44^. This step identified organs where the protein set is significantly overrepresented. Enrichment was computed using GSEA-based scoring. Only medium and high expression levels were considered. The significance of each enrichment score was evaluated using permutation testing, where gene labels were randomly shuffled (10,000 permutations), and empirical *p*-values were calculated as the proportion of permuted scores exceeding the observed enrichment score. Holm-adjusted *p*-values were used for multiple testing correction. Tissues with adjusted *p*-values < 0.05 were retained for downstream single-cell resolution analysis. Next, organs identified as significantly enriched in the previous step were further analyzed using single-cell RNA-sequencing data from the Tabula Sapiens atlas ^45^. For each organ, enrichment was computed across all annotated cell types using log normalized transcriptomic expression levels. Expression thresholds (log normalized expression > 1) were used to define expressed genes. To interpret the functional significance of the results, leading edge subsets were extracted from the single-cell analysis and a pathway enrichment analysis was done in Reactome (v94) ^46^, with significantly (FDR < 0.05) enriched modules reported.

## Results

### Study design and initial unsupervised characterization of the GNPC plasma proteome dataset

To establish a robust analytical foundation, we began with Version 1 of the GNPC dataset, which comprises plasma proteomic profiles generated across 16 international contributing sites, totalling 13,733 individuals across five major neurodegenerative diseases: AD (*n* = 4,217; 13 sites), FTD (*n* = 175; 6 sites), PDD (*n* = 185; 10 sites), PD (*n* = 540; 8 sites), and ALS (*n* = 244; 1 site), alongside non-impaired (NI) controls (*n* = 8,372; 15 sites) (Fig. 2a; Supplementary Table 1) ^17^. Proteomic quantification was performed using the SomaScan 7k platform, yielding 6,340 unique plasma protein measurements per participant.

**Figure 1.**
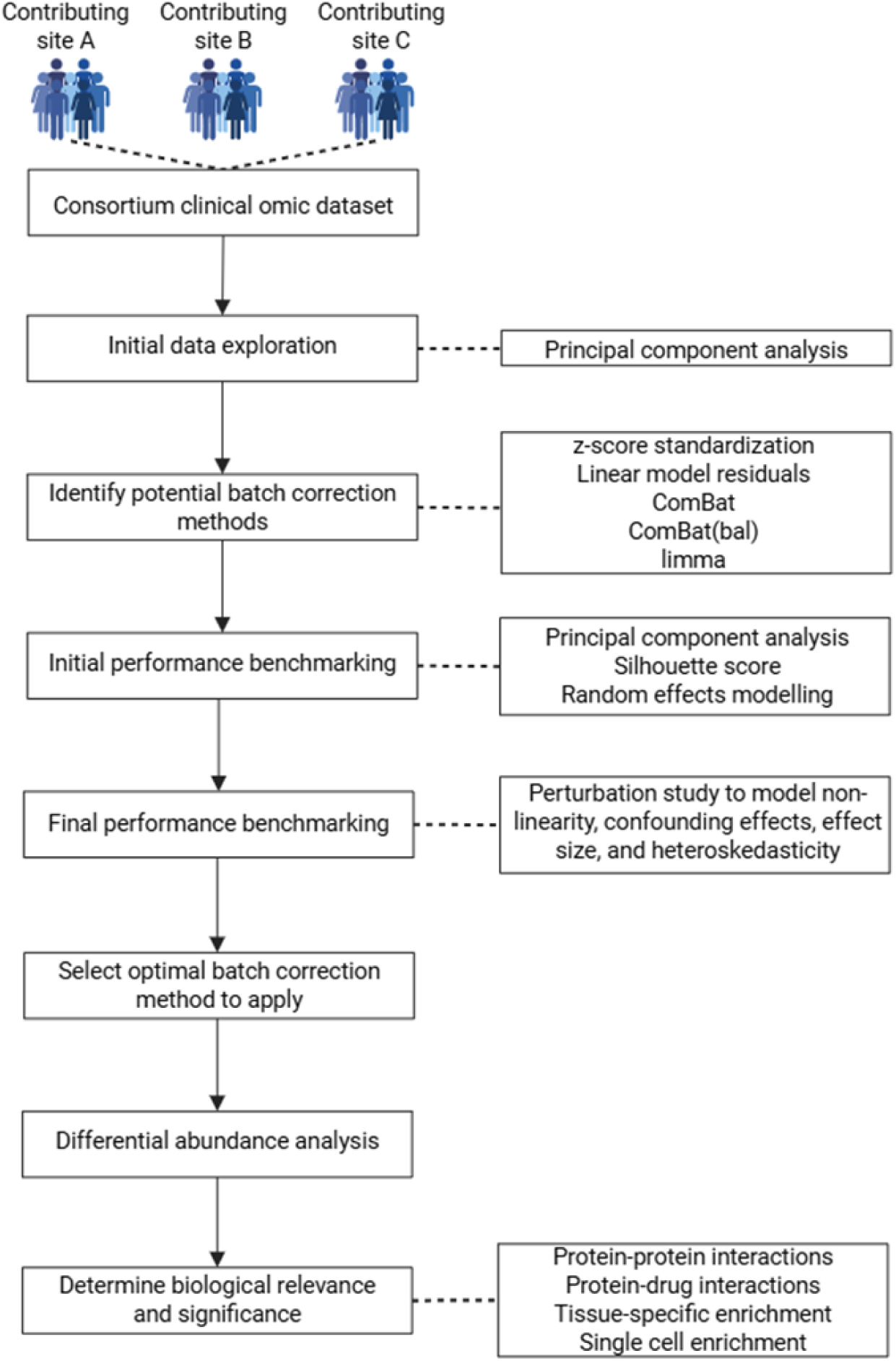
Methodological pipeline of the current study and for analyzing consortium-generated clinical omic datasets. Clinical omic data is collected from across various contributing clinical sites. Data is then initially explored using a principal component analysis (PCA). Potential batch correction methods are identified and their performance benchmarked using methods including PCA, silhouette score, and random effects modelling. The performance of batch correction methods is then benchmarked via a perturbation study to model non-linearity, confounding effects, effect size, and heteroskedasticity. Based on these benchmarking metrics, an optimal batch correction method is selected, and differential abundance analysis is done to identify molecular candidates. The biological relevance and significance of the identified molecular candidates is assessed using methods including protein-protein interactions, protein-drug interactions, and tissue-specific and single cell enrichments.

**Figure 2.**
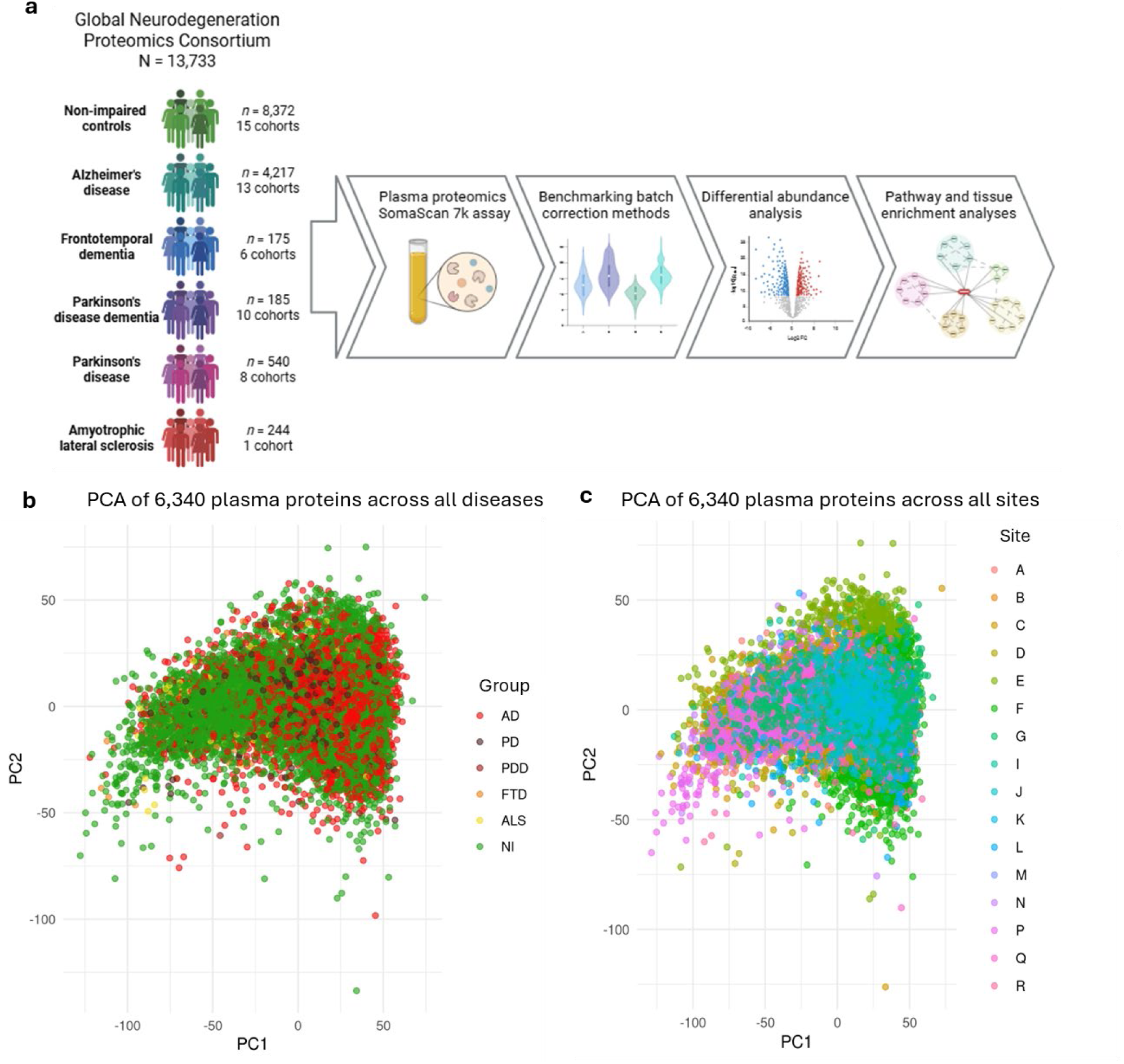
Study design and principal component analyses (PCA) of the raw GNPC cohort plasma proteomic dataset. (**a**) Overall design of the current study. (**b**) PCA of 6,340 plasma proteins measured in participants with five different neurodegenerative diseases and non-impaired controls showing little distinctive clustering based on disease. (**c**) PCA of 6,340 plasma proteins measured in participants from 16 different contributing sites showing little distinctive clustering based on site. Abbreviations: AD: Alzheimer’s disease; ALS: amyotrophic lateral sclerosis; FTD: frontotemporal dementia; NI: non-impaired controls; PCA: principal component analysis; PD: Parkinson’s disease; PDD: Parkinson’s disease dementia.

Initial unsupervised visualization using principal component analysis (PCA) revealed minimal separation among diagnostic groups (Fig. 2b) and similarly limited stratification when samples were delineated by contributing site (Fig. 2c). However, the apparent absence of discrete clusters in PCA does not indicate a lack of structured heterogeneity. PCA is intrinsically insensitive to high-cardinality batch variables and often compresses multi-layered confounding into diffuse low-variance components, particularly in datasets of this scale and imbalance (13,733 participants spanning 16 contributing sites across multiple diagnoses) ^47^. Under such conditions, subtle but systematic site-level effects are unlikely to manifest as distinct clusters yet can still exert pronounced influence on downstream association analyses, classification performance, and biomarker discovery ^26,27,48,49^.

These observations highlight a central analytical challenge for consortium-scale proteomics: the absence of visible clustering in low-dimensional projections is not sufficient to rule out structured technical heterogeneity. In large, multi-site datasets where clinical and technical factors are often partially confounded, even subtle cross-site differences may influence downstream analyses. At the same time, uniform or overly aggressive correction approaches risk suppressing biologically meaningful variation, particularly in smaller or clinically distinct subgroups. Accordingly, before drawing biological conclusions or conducting differential analyses, it is essential to formally quantify the relative contributions of site, diagnosis, and participant-level covariates to proteomic variance.

### Systematic decomposition of variance reveals the need for explicit modelling of site heterogeneity

We therefore next sought to systematically decompose variance across diagnosis, site, and participant-level factors to determine whether, and to what extent, site heterogeneity must be explicitly modelled to enable valid inference. To do so, we focused on Alzheimer’s disease (AD), the diagnostic group with the greatest number of contributing sites (*n* = 13) and participants (*n* = 4,217), providing a rigorous test case for assessing variance structure and correction strategies under conditions of maximal site heterogeneity. Consistent with our initial observations in the full cohort, PCA of AD versus NI controls across all 6,340 plasma proteins showed no clear diagnostic separation (Fig. 3a), and sample delineation by site similarly did not produce distinct clusters (Fig. 3b). However, mixed-effects variance partitioning of the uncorrected data revealed that site accounted for 7.9% of total proteomic variance, whereas diagnostic group explained <1% (Fig. 3c-d). This imbalance indicates that technical heterogeneity substantially exceeds disease-associated signal in raw plasma proteomics. This is consistent with prior observations in different tissues and platforms ^50^ and underscores the need for rigorous harmonization prior to biological interpretation.

**Figure 3.**
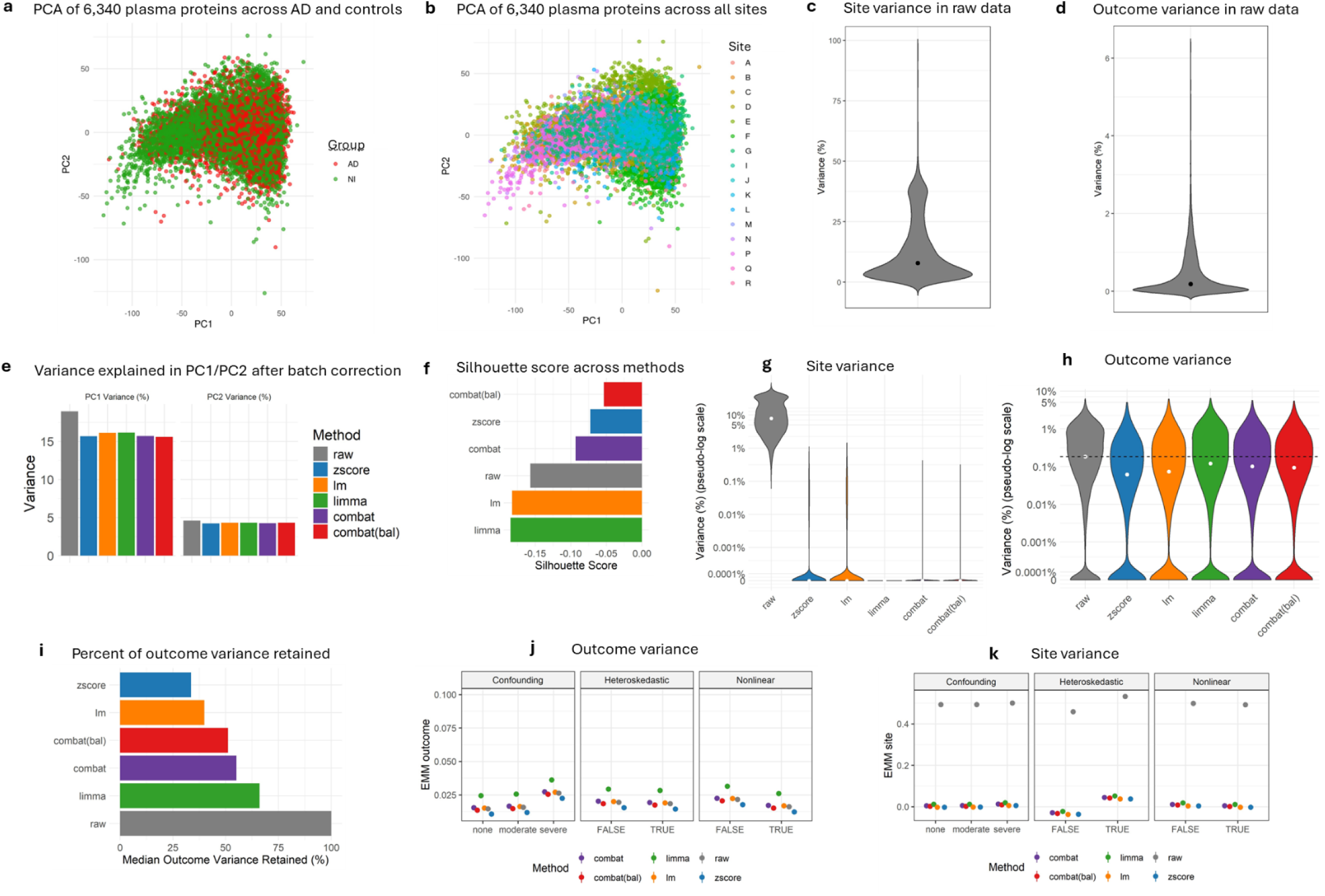
Batch effects are present in the raw plasma proteomic data after integrating across multiple sites and these are differentially affected by batch correction methods. (**a**) Principal component analysis (PCA) of 6,340 plasma proteins measured in patients with AD and NI controls showing no disease-specific group clustering. (**b**) PCA of 6,340 plasma proteins measured in all individuals from across 16 contributing sites similarly revealing no site-specific clustering of cases. (**c**) Random effects modelling showing 7.9% of the per-protein variance is attributable to site in the plasma proteomic dataset. (**d**) Random effects modelling showing that less than 1% of the per-protein variance is attributable to outcome (diagnosis) in the plasma proteomic dataset. (**e**) Variance explained by principal component (PC) 1 and PC2 in the raw plasma proteomic dataset and following batch correction using z-score, linear model (LM) residuals, ComBat, ComBat(bal), and limma. All methods reduced PC1 variance to 15-16% whereas PC2 variance was maintained at around 4%. (**f**) Silhouette score using site labels across the top 10 PCs showing reduced site-based clustering across batch correction methods. (**g**) Random effects modelling showing that all five batch correction methods eliminate site-related variance. ComBat, ComBat(bal), and limma also show minimal residual tails, suggesting fewer proteins remain influenced by site after correction. (**h**) Random effects modelling showed that LM residuals and z-score suppressed outcome-related (biological) signal to a greater degree than ComBat, ComBat(bal), and limma. (**i**) Proportion of outcome variance retained by the correction methods showing limma retains the greatest proportion. (**j**) Outcome-related variance across the perturbation study showing that limma consistently preserves the highest level of outcome-related variance under different analytical conditions. (**k**) Site-related variance across the perturbation study showing that all batch correction methods effectively remove site-related variance across different conditions.

To evaluate how standard correction approaches reshape this variance architecture, we benchmarked five widely used strategies, z-score normalization, linear model (LM) residualization, ComBat, ComBat with site-rebalancing (ComBat(bal)), and limma. All methods reduced PC1 variance from 21% (raw) to ∼15-16% post-correction, while PC2 remained stable at ∼4% (Fig. 3e). Silhouette widths computed across the top ten PCs revealed partial but heterogeneous improvements in site mixing: ComBat, ComBat(bal), and z-score shifted site-wise silhouette scores toward zero (-0.05 to -0.09), whereas LM residuals and limma produced more negative scores (∼-0.18), reflecting stronger disruption of site-structured similarity (Fig. 3f). Given that silhouette width is descriptive and sensitive to global covariance structure ^24^, we next directly quantified residual variance attributable to site and diagnosis following correction. All methods achieved near-complete elimination of site-associated variance (Fig. 3g). However, their effects on biological signal differed markedly.

LM residuals and z-score normalization suppressed disease-related variance most severely, whereas ComBat, ComBat(bal), and limma retained a larger proportion of outcome-associated signal (Fig. 3h). Limma preserved the greatest fraction of diagnostic variance (66%), compared to ComBat and ComBat(bal), with z-score retaining the least (33.6%) (Fig. 3i). These findings indicate that although multiple strategies can remove technical site effects, model-based empirical approaches provide superior balance between technical harmonization and preservation of disease-relevant signal.

To establish the robustness of these findings, we conducted a large-scale perturbation analysis in which 2,000-subject, 1,000-protein subsets were repeatedly sampled (*n* = 100) under systematically varied conditions, including controlled batch-outcome confounding (none, moderate, severe), heteroskedasticity, nonlinear effects, and variable signal sparsity. Across nearly all perturbations, limma consistently preserved the highest level of outcome-related variance while achieving near-complete elimination of site-associated variance, matching or exceeding the performance of ComBat and ComBat(bal) (Fig. 3j-k).

Together, these analyses demonstrate that site effects account for substantially more variance than diagnosis in raw plasma proteomic data and must be explicitly modelled for valid biological inference. While several approaches effectively suppress site-related variance, model-based harmonization, particularly limma, achieves the best balance of removing technical heterogeneity while preserving disease-relevant biology. These results establish limma as the best performing batch correction method for multi-site plasma proteomic integration and form the analytical foundation for downstream mechanistic and biomarker discovery analyses in this study.

### Shared vulnerability pathways and disease-specific molecular dysfunction across neurodegenerative diseases

To identify plasma proteins that reflect conserved biological processes underlying neurodegenerative disease, rather than site-specific or technical artefacts, we implemented a two-pronged differential analysis framework. First, we applied a limma-based linear model across all contributing sites to detect diagnosis-associated proteins in the harmonized dataset. Second, we performed site-stratified differential expression analyses (restricted to sites with >10 cases and >10 controls) followed by inverse-variance weighted fixed-effects meta-analysis. To distinguish reproducible biological signal from signatures driven by site composition or sampling imbalance, we introduced three meta-analytic diagnostics: (1) maximum site weight (>60%), identifying cases where one site disproportionately drives the pooled estimate; (2) opposing weight (>30%), quantifying conflicting directional effects; and (3) leave-one-site-out (LOSO) fragility, identifying proteins whose direction or significance is not stable to exclusion of any single site (Extended Data Fig. 1a-b). Integrating pooled estimates, meta-analytic results, and diagnostic metrics, we classified proteins into five confidence tiers (Methods Table 1; Extended Data Fig. 2a-b).

Using this framework, we identified *n* = 2,524 AD, 580 PD, 101 FTD, 85 ALS, and 64 PDD high-confidence plasma protein associations relative to NI controls (Supplementary Tables 2-6). Functional enrichment analyses revealed a mixture of shared and disease-specific biology (Fig. 4a-c; Supplementary Tables 7-9). A prominent core signature was shared between AD and PD, involving cell stress adaptation and immune-metabolic regulation, including negative regulation of apoptosis, glycolytic and gluconeogenic metabolism, viral response, protein phosphorylation, angiogenesis, blood coagulation, and cell adhesion (Fig. 4a; Supplementary Table 7). These convergent processes suggest coordinated disruption of metabolic-immune coupling and tissue remodelling in both diseases. Despite this common axis, AD exhibited the broadest systemic molecular footprint, with additional enrichment in intracellular trafficking, membrane receptor signalling, and metabolic control (Fig. 4a; Supplementary Table 7). PD showed a more focused enrichment, dominated by ubiquitin-mediated proteostasis (Fig. 4a; Supplementary Table 7). PANTHER molecular function analysis reinforced these distinctions. AD and PD both demonstrated enrichment in kinase, receptor, and cytokine/chemokine binding activities, indicating shared disruption of phospho-signaling and inflammatory mediator networks (Fig. 4b; Supplementary Table 8). In contrast, PD-specific enrichment of ubiquitin-protein transferase activity highlights proteostatic failure as a core mechanistic feature of PD (Fig. 4b; Supplementary Table 8). These patterns extended to KEGG pathways. AD was characterized by systemic pathway disruption, with broad enrichment across complement activation, phagocytic signalling, metabolic reprogramming, extracellular matrix remodelling, and second messenger signalling (Fig. 4c-h; Supplementary Table 9). PD mirrored many of these pathways but uniquely retained ubiquitin-proteasome and T cell receptor signalling, further emphasizing proteostasis and adaptive immune signalling imbalance (Fig. 4c-g; Supplementary Table 9).

**Figure 4.**
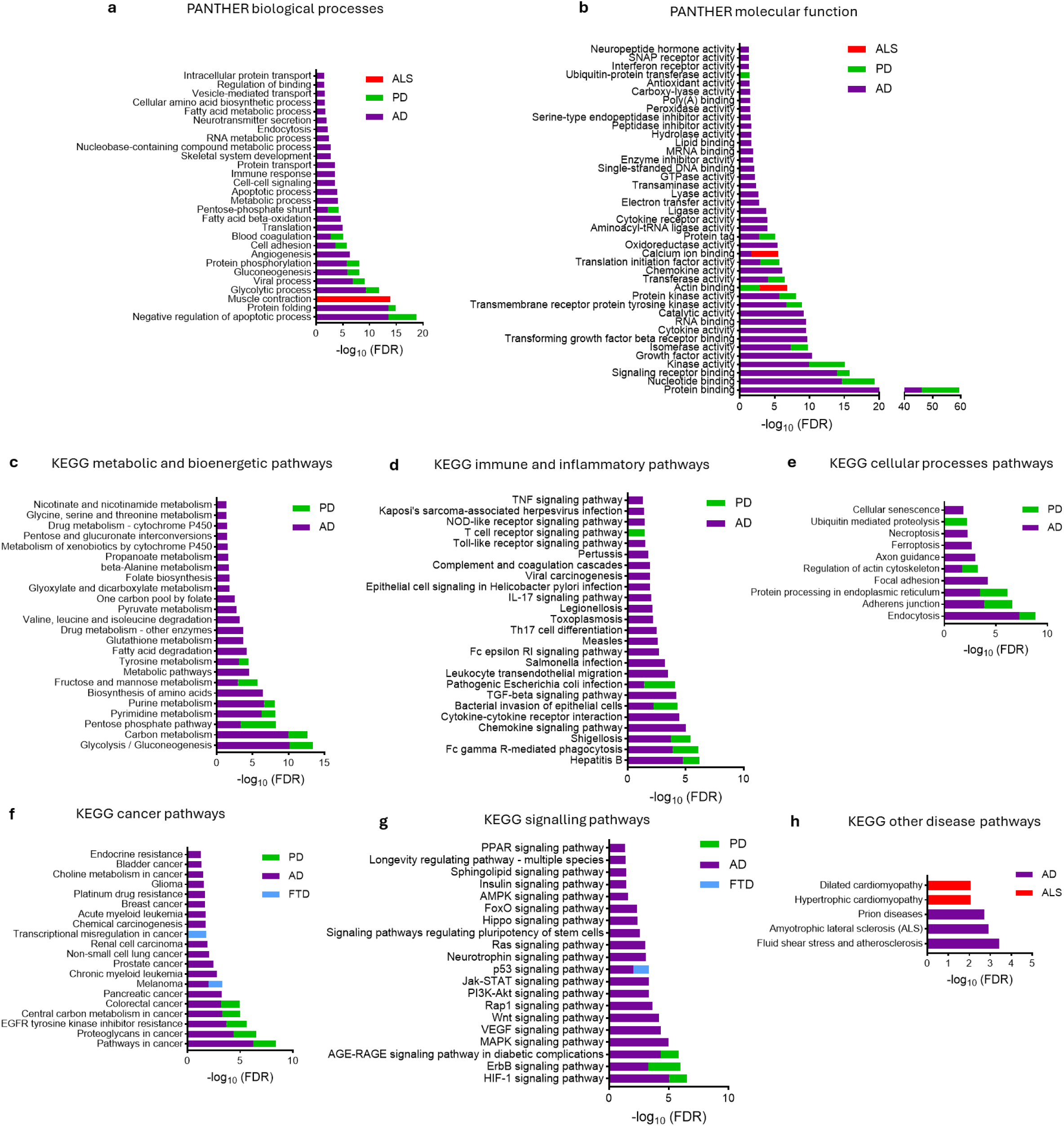
Functional enrichment analyses of exploratory plasma proteins across neurodegenerative diseases. (**a**) Significantly enriched PANTHER biological processes in AD, PD, and ALS plasma proteins. (**b**) Significantly enriched PANTHER molecular functions in AD, PD, and ALS plasma proteins. (**c**-**h**) Significantly enriched KEGG (**c**) metabolic and bioenergetic, (**d**) immune and inflammatory, (**e**) cellular processes, (**f**) cancer, (**g**) signalling, and (**h**) other disease pathways in AD, PD, ALS, and FTD plasma proteins. Abbreviations: ALS: amyotrophic lateral sclerosis; AD: Alzheimer’s disease; FTD: frontotemporal dementia; PD: Parkinson’s disease.

By contrast, ALS exhibited a markedly narrower systemic plasma signature. Biological processes enrichment was confined to muscle contraction (Fig. 4a; Supplementary Table 7), and molecular functions were restricted to actin and calcium ion binding, consistent with changes in cytoskeletal integrity and excitation-contraction coupling (Fig. 4b; Supplementary Table 8). KEGG enrichment in cardiomyopathy-associated pathways further supported peripheral neuromuscular involvement (Fig. 4h; Supplementary Table 9). FTD showed limited but biologically relevant enrichment, overlapping with AD in p53 and JAK–STAT signalling and uniquely implicating transcriptional misregulation (Fig. 4f-g; Supplementary Table 9). PDD did not show statistically significant enrichment, consistent with limited sample size and clinical heterogeneity rather than absence of biological changes.

These findings demonstrate that neurodegenerative diseases share a common axis of immune-metabolic stress and tissue remodelling. However, disease-specific biological dysfunction differentiates their pathological profiles: proteostasis disruption in PD, transcriptional dysregulation in FTD, and cytoskeletal and excitation-contraction imbalance in ALS, with AD exhibiting the most extensive systemic involvement. These shared and disease-specific biological signatures provide defined targets for dissecting disease mechanisms and developing mechanistically informed therapeutic strategies.

### Immune and endocrine remodelling and mitochondrial redox stress define AD-specific systemic signatures and druggable network hubs

To extend mechanistic interpretation beyond pathway-level annotations, we next assessed the organ and cellular origins of disease-associated plasma proteins. Consistent with the limited pathway enrichment observed in ALS, FTD, and PDD, none of these disorders exhibited significant organ-level enrichment, likely reflecting reduced cohort size and increased clinical and molecular heterogeneity relative to AD and PD. In PD, disease-associated proteins showed significant enrichment in lymphoid organs, including lymph nodes and bone marrow (Fig. 5a; Supplementary Table 10). However, cell type-specific analyses revealed no preferential enrichment of discrete immune subsets, suggesting diffuse immune and hematopoietic involvement rather than activation of a restricted cell types.

**Figure 5.**
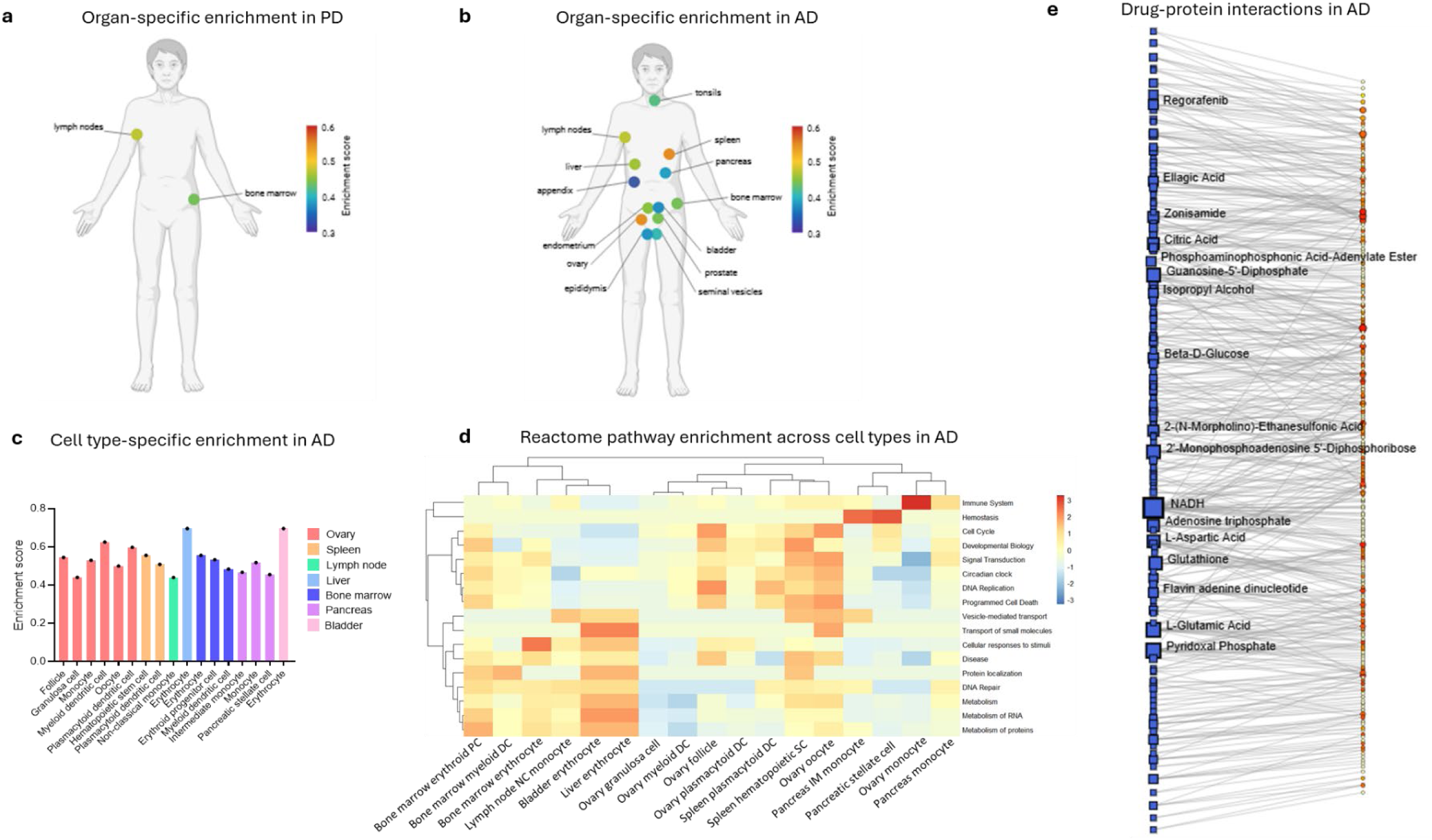
Organ, cell-specific, and druggable network enrichment analysis of exploratory plasma proteins across AD and PD. (**a**-**b**) Organ-specific enrichment in (**a**) PD and (**b**) AD. Coloured scale bar represents enrichment score. (**c**) Cell type-specific enrichment across organs implicated by AD plasma proteins. (**d**) Reactome pathway enrichment of leading-edge nodes across specific cell types in AD. Scale bar represents -log10(FDR). (**e**) Drug-protein interaction network analysis across AD plasma proteins. Blue squares represent pharmacologically active compounds, with the size of the square representative of the degree and betweenness. Orange circles represent the >2,000 AD plasma proteins identified in the study. Named pharmacologically active compounds are those with at least seven protein interactions.

In contrast, AD plasma proteins demonstrated widespread and highly significant enrichment across multiple organ systems, including immune and lymphoid tissues (tonsil, lymph node, spleen, appendix, bone marrow), metabolic organs (liver, pancreas), reproductive tissues (ovary, endometrium, prostate, seminal vesicle, epididymis), and the urogenital tract (bladder) (Fig. 5b; Supplementary Table 10). The strongest enrichment occurred in the spleen and ovary, indicating that systemic immune remodelling and reproductive-hormonal axis dysregulation represent major contributors to the AD-associated circulating proteome. Cell type-specific analyses further revealed that ovary, bone marrow, and pancreas harboured the greatest diversity of enriched cellular populations (Fig. 5c; Supplementary Table 11). Notably, monocytes, erythroid lineage cells, and dendritic cell subsets were recurrently enriched across multiple organs, indicating a coordinated, multi-organ immune-hematopoietic activation state, rather than isolated local organ involvement. Leading-edge gene set integration across enriched organs and cell types revealed three coherent biological modules (Fig. 5d; Supplementary Table 12). First, an innate immune-hemostatic activation module, driven largely by tissue-resident monocytes and stromal/stellate cell populations in pancreas and ovary, consistent with peripheral inflammatory priming and microvascular remodelling (Fig. 5d). Second, a proliferative-developmental regulation module, reflecting cell-cycle progression, chromatin and transcriptional remodelling, and developmental lineage reactivation, most pronounced in granulosa cells, hematopoietic progenitors, oocytes, and plasmacytoid dendritic cells (Fig. 5d). Finally, an anabolic translational and metabolic efficiency module, enriched in erythroid and erythropoietic lineages, indicative of elevated ribosomal load, oxidative metabolic demand, and proteostasis burden (Fig. 5d).

To determine whether the proteomic signatures are pharmacologically tractable, we performed drug-protein interaction network analysis on the AD-associated plasma proteins. High-centrality compounds were strongly enriched for redox cofactors and electron transport mediators (NADH, FAD, glutathione, pyridoxal phosphate), indicating disrupted redox homeostasis and mitochondrial bioenergetics as core AD-specific metabolic targets (Fig. 5e; Supplementary Table 13). Central carbon metabolites (β-D-glucose, citric acid, ATP, L-glutamate, L-aspartate) further linked AD to altered glycolytic-TCA coupling and neurotransmitter-linked anaplerotic metabolism (Fig. 5e; Supplementary Table 13).

Compounds associated with post-translational modification and proteostasis control (GDP cycling, ADP-ribosylation intermediates, adenylate analogs) mapped to broad intracellular stress adaptation and protein-quality control systems (Fig. 5e; Supplementary Table 13).

Finally, the presence of zonisamide and regorafenib highlighted pharmacologically accessible intersections within ion handling and kinase-regulated signalling, respectively, both of which were prominently represented in the AD plasma network.

Together, these results show that AD is characterized by system-wide immune and stromal activation coupled with impaired mitochondrial redox balance, pointing to metabolic-oxidative regulation and peripheral immune signalling as key determinants of systemic AD biology.

### Prioritized disease-specific plasma biomarkers for clinical translation in neurodegeneration

To identify potential plasma biomarkers, we implemented a secondary, conservative analysis designed to prioritize demographically robust and disease-specific protein associations. For each diagnostic group, we performed 1:1 case-control matching by age and sex against the NI reference pool, minimizing demographic confounding and yielding biomarkers that are likely to be highly disease-specific. The AD cohort was already age and sex balanced relative to NI controls; therefore, the AD high confidence protein set was retained in full under the matched analyses (Supplementary Table 2). Among the top 15 prioritized AD proteins, 11 were specific to AD and did not appear as high confidence markers in any other neurodegenerative disease. These included components of lipoprotein metabolism (APOB), complement and inflammatory signalling (C3, GDF2), nucleotide and redox homeostasis (CTPS1, NT5C, GLRX3), and protein folding and translational control (CCT5, PA2G4, RRS12, ALAD, PGP), together indicating systemic metabolic-immune dysregulation as a core AD plasma signature (Fig. 6; Supplementary Table 2). In contrast, FTD yielded only a single robust, disease-specific candidate, HS6ST3. This is a regulator of heparan sulfate biosynthesis and extracellular signalling (Fig. 6; Supplementary Table 14), consistent with the limited systemic proteomic signal and greater clinico-pathological heterogeneity observed in FTD. No protein associations survived the matched analysis in PDD (Supplementary Table 15), reflective of the underpowered nature of PDD cases. In PD, 12 proteins remained robust after matching (Supplementary Table 16). However, only SUMF1 (sulfuric ester metabolism) and TPPP3 (tubulin and microtubule assembly) were PD-specific. (Fig. 6). By contrast, ALS displayed a distinct and highly specific plasma profile, with 10 of the top 15 candidates unique to ALS. These included proteins associated with sarcomere structure (MYOM2, MYBPC1, ACTN2, TNNI2), muscle regeneration and contractile machinery (KLHL41, PDLIM3, APOBEC2), and neuromuscular synaptic organization (LRTM1, ART3) (Fig. 6; Supplementary Table 17), consistent with peripheral neuromuscular involvement as a defining systemic feature of ALS.

**Figure 6.**
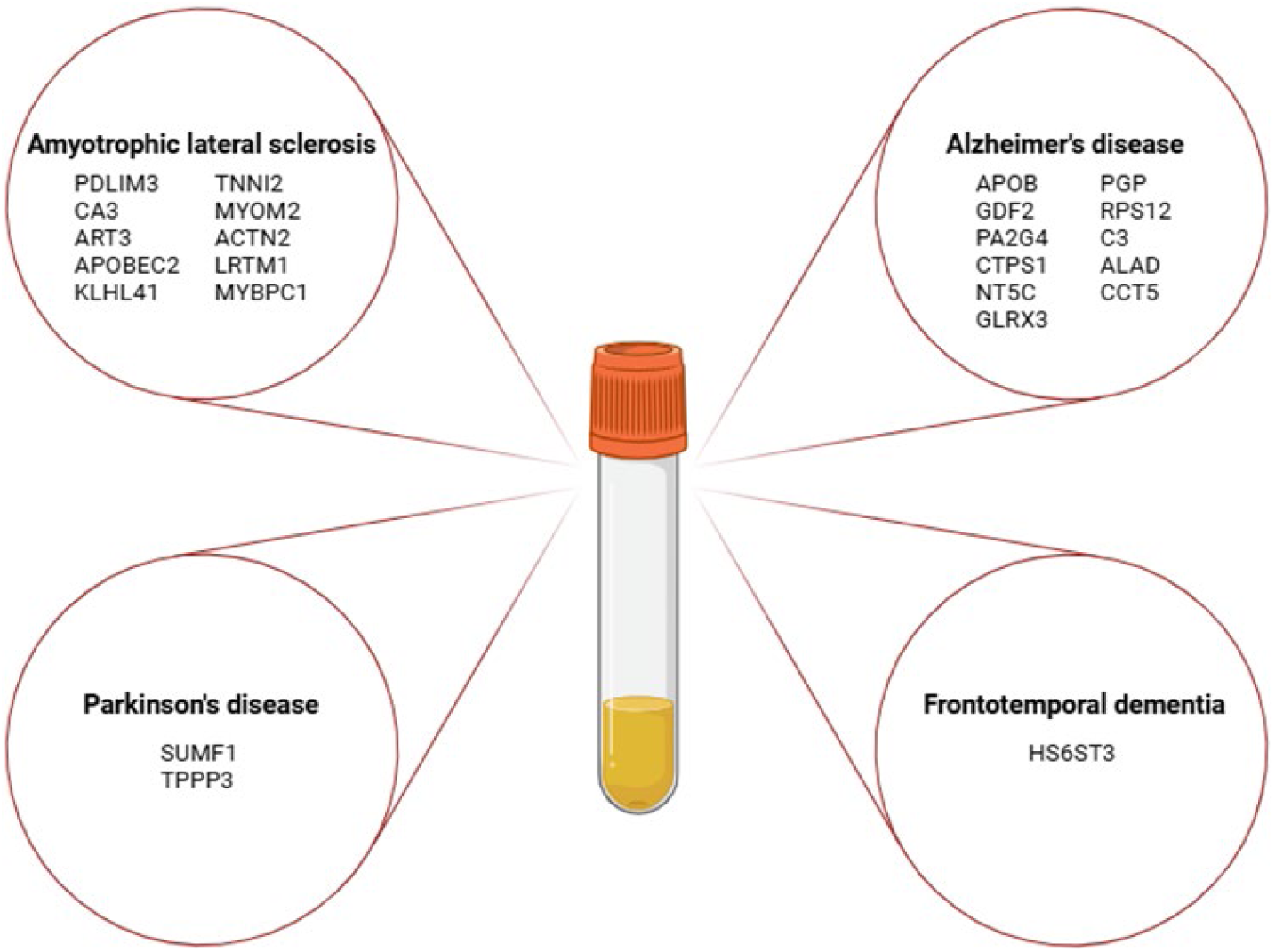
Identified candidate plasma protein biomarkers for neurodegenerative diseases. The listed proteins represent those that are high confidence candidate plasma proteins and specific to a single neurodegenerative disease.

Overall, these analyses identify disease-specific plasma protein candidates with high translational robustness, providing a focused set of prioritized biomarkers for clinical validation and stratification efforts across neurodegenerative diseases.

## Discussion

Our study demonstrates how large scale, multi-center plasma proteomics can substantially refine the molecular understanding of neurodegenerative diseases. By harmonizing proteomic data from over 13,000 individuals across 16 international contributing sites in the GNPC, we established a next-generation framework to investigate both shared and disease-specific biology across AD, PD, FTD, PDD, and ALS. This unprecedented scale, diversity, and standardization enabled the detection of subtle yet reproducible protein signatures that would likely remain elusive in underpowered single-center studies. Indeed, these strategic analyses mirror the transformative impact of other coordinated efforts in biomedicine, such as CPTAC and AMP consortiums, underscoring that integrated, multi-site datasets are essential to disentangle the systemic complexity of human disease ^5,7,10,17^. By demonstrating that the circulating proteome harbors clinically meaningful signatures of disease pathophysiology, our findings position plasma proteomics as a scalable and accessible platform for mechanistic discovery, biomarkers for patient stratification, and identification of potential therapeutic strategies.

Our results also highlight a central methodological challenge in consortium-scale proteomics: technical heterogeneity can easily rival or exceed biological variability. In the GNPC dataset, we observed that variance attributable to contributing site far exceeded variance due to diagnosis, consistent with prior evidence that differences in sample handling, assay calibration, and site-specific protocols introduce structured batch effects that can spuriously inflate associations and undermine reproducibility ^23–26^. However, our benchmarking showed that the solution is not simply to maximally suppress technical factors. Overly aggressive harmonization (e.g. global normalization or naive linear residualization) can inadvertently remove biologically meaningful signal, especially when technical and clinical variables are confounded. In contrast, a model-based empirical approach, specifically using limma, achieved a principled balance, eliminating site-related artifacts while preserving the majority of disease-related signals across a range of confounded and heteroskedastic scenarios. These findings underscore that analysis choices fundamentally shape biological interpretation, and that rigorous, data-driven harmonization is essential for reproducible mechanistic and biomarker discovery. They also emphasize the need for data-generating consortia to accompany resource releases with consensus harmonization guidelines based on thorough benchmarking. Without such guidance, analytic heterogeneity could become a dominant driver of inference, fragmenting biological insights across studies and hindering reproducibility.

Leveraging our harmonized dataset, we uncovered a prominent convergent axis of immune–metabolic dysregulation spanning AD and PD. Both AD and PD shared a plasma signature of cellular stress and metabolic-immune disruption, with pathways related to apoptotic regulation, glycolytic and gluconeogenic metabolism, inflammatory signalling, angiogenesis, and tissue remodelling. This convergence suggests a coordinated breakdown of peripheral metabolic-immune homeostasis in neurodegeneration, aligning with extensive evidence that chronic systemic inflammation, mitochondrial dysfunction, and metabolic stress contribute to AD and PD pathogenesis ^51–58^. Despite this shared axis, AD and PD diverged markedly in the breadth and composition of their proteomic profiles. AD exhibited the most extensive systemic involvement. We observed widespread enrichment in complement activation, stromal-immune signalling, metabolic reprogramming, extracellular matrix remodelling, and second messenger pathways. This expansive footprint reinforces the view of AD as a multi-organ, multi-system disorder rather than a purely neurocentric pathology ^59–61^. Yet most mechanistic and biomarker studies remain brain-centred, with relatively little attention to early peripheral changes, particularly in prodromal disease. Our findings argue for a concerted shift towards characterising these peripheral and system-wide alterations, which are likely to arise years before overt neurodegeneration, to better understand disease initiation, progression, and opportunities for intervention. By contrast, PD’s plasma profile was more circumscribed, centered on ubiquitin-mediated proteostasis and adaptive immune signalling, with strong enrichment of the ubiquitin-proteasome system and T-cell receptor signalling. These findings cement proteostatic failure and immune-synaptic dysregulation as core mechanistic hallmarks of PD ^62–66^.

Beyond AD and PD, the other neurodegenerative disorders showed more restricted but biologically distinct systemic proteomic changes. ALS was defined by a strikingly specific plasma signature dominated by proteins involved in skeletal muscle structure and neuromuscular function. Many of the top ALS-associated proteins were structural muscle components (e.g. MYBPC1), muscle development/regeneration factors (e.g. KLHL41), or synaptic organizers at the neuromuscular junction (e.g. LRTM1). This peripheral muscle-derived signature reflects the fundamental motor neuron degeneration and muscle denervation in ALS, and indeed we found enrichment of pathways related to cytoskeletal remodelling and excitation-contraction coupling. Such findings are consistent with the concept that skeletal muscle degeneration and its metabolic sequelae are detectable in circulation ^67–70^. Notably, our results parallel prior reports of broad increases in muscle-contractile and extracellular matrix proteins in ALS blood ^17^. The clear delineation of ALS by a muscle protein signature underscores that ALS, while a primary neurodegenerative disease, is also a multi-system disorder with peripheral involvement, and it provides a focused panel of translational biomarker candidates unique to ALS.

FTD, in contrast, showed only very modest plasma proteomic changes. FTD yielded a small but coherent signature implicating dysregulated transcriptional stress responses, most prominently involving p53 and JAK-STAT signalling. Although there is limited genetic data available in the GNPC dataset, this finding in alignment with evidence linking *C9orf72*-associated FTD to p53-mediated neuronal vulnerability and JAK-STAT hyperactivation ^71,72^. PDD did not show any significant plasma protein enrichment in our analysis, likely due to the limited cohort size and clinical heterogeneity of the PDD group ^73^ rather than a true absence of systemic changes. This lack of signal highlights the need for larger, dedicated studies of PDD to determine whether distinct peripheral biomarkers can be identified for that subtype.

To further interpret the tissue and cellular origins of these disease signatures, we performed organ- and cell-enrichment analyses. This revealed striking disease-specific patterns of peripheral involvement. PD-associated proteins were enriched for those originating from lymphoid organs, pointing to diffuse hematopoietic immune activation rather than activation of a single immune cell subset ^74–76^. By contrast, AD-associated proteins mapped to a broad array of tissues, notably including immune-related organs (such as spleen) and endocrine/metabolic organs (such as ovary), suggesting coordinated immune activation ^77,78^ along with dysregulation of the reproductive hormonal axis in AD ^79–81^. At the cellular level, we found recurring contributions from monocytes, erythroid lineage cells, and dendritic cells across multiple enriched organs. These convergent cell signals point to at least three underlying systemic modules common to the diseases. One module involves innate immune activation coupled with coagulatory and microvascular remodelling processes ^77,82,83^. A second module reflects proliferative and chromatin-remodelling, suggesting reactivation of developmental or stress-response pathways ^84,85^. A third module involves enhanced metabolic and anabolic activity in erythroid cells, indicative of altered erythropoietic function ^86–88^.

Importantly, a network-based pharmacologic analysis highlighted several druggable nodes within the AD plasma signature. These included redox cofactors, electron transport intermediates, central carbon metabolism regulators, and proteostasis factors, indicating that mitochondrial redox balance and metabolic-oxidative coupling are actionable therapeutic entry points in AD ^89–91^.

Using demographically matched case-control analyses, we next identified specific circulating proteins that were significantly altered in each disease, highlighting candidates for blood-based biomarkers. In AD, the top plasma biomarkers converged on the same metabolic and immune pathways noted above. Notably, apolipoprotein B (APOB) was elevated in AD plasma, reinforcing epidemiological links between mid-life dyslipidemia and AD risk ^92–94^.

Consistent with this, prior studies have found higher APOB in the blood of AD patients and even its presence in CSF of individuals at risk ^92,93^, suggesting that peripheral lipid metabolism contributes to AD pathology. We also observed increased complement C3 in AD, in line with the known involvement of innate immunity and complement activation in AD ^52,77,95^. Interestingly, many of these plasma markers differ from the classical AD biomarkers (amyloid-β and tau), implying that proteins such as APOB, GDF2, and the nucleotide metabolism enzymes CTPS1 and NT5C reflect parallel systemic processes that complement the central hallmarks of AD. One factor likely contributing to this pro-inflammatory, metabolic signature is *APOE* genotype. Recent GNPC work showed that *APOE* ε4 carriers have a heightened inflammatory proteomic profile ^96^, and since *APOE* ε4 is overrepresented in AD cohorts, this genotype may partially drive the observed plasma differences. Overall, our findings underscore systemic metabolic-immune dysregulation as a core feature of AD, echoing the broader literature linking mid-life metabolic syndrome and chronic inflammation to elevated AD risk. As these candidate markers are readily measurable in blood, they hold promise to augment traditional CSF and imaging biomarkers in tracking AD progression or therapeutic responses in clinical trials.

In contrast to AD, FTD and PD yielded far fewer robust plasma biomarkers under the same stringent criteria. FTD showed a striking paucity of significant changes, with only a single protein, HS6ST3, emerging as a candidate biomarker. HS6ST3 encodes a heparan sulfate sulfotransferase, pointing toward altered heparan sulfate biology in FTD. This biomarker was also identified in the initial GNPC study ^17^, and it aligns with evidence that heparan sulfate proteoglycans co-deposit with pathological tau and modulate its aggregation in tauopathies ^97^. However, it is unclear if this finding is driven by a particular etiology of FTD, such as FTLD-TDP or FTLD-tau, as the consortium FTD diagnoses available are not that specific. Future research would benefit from examining this potential biomarker in distinct FTD groups. PD yielded an intermediate number of candidates (12 after matching), but only two were unique to PD: SUMF1 and TPPP3. SUMF1 (sulfatase-modifying factor 1) is a facilitator of lysosomal enzyme function, implicating lysosomal and proteostatic dysfunction in PD ^17^. TPPP3 (tubulin polymerization-promoting protein 3) is a microtubule-associated protein involved in cytoskeletal assembly and has been linked to α-synuclein aggregation ^98,99^. Notably, TPPP3 dysregulation has also been observed in longitudinal studies of prodromal PD, suggesting that cytoskeletal network changes precede clinical motor onset ^100^. Consistent with clinical overlap, the PDD group did not yield any exclusive biomarkers; this is likely because PDD shares much of its proteomic profile with PD and/or AD ^101^ and was underpowered in our analysis. Expanding the representation of PDD in future cohorts will be important to determine if any unique plasma markers characterize that condition. Overall, our PD results reinforce the importance of lysosomal dysfunction and cytoskeletal alterations in PD pathogenesis, and they provide two concrete plasma candidates for further validation as PD-specific biomarkers. By comparison, as noted above, the plasma markers distinguishing ALS were almost entirely muscle-derived, underlining the peripherally driven profile unique to ALS.

Despite the strengths of our multi-site proteomic approach, our study has certain limitations. First, the cross-sectional design captures snapshots of the disease proteome; longitudinal studies will be needed to track how these protein signatures evolve with disease progression or in response to therapy. Second, while the SomaScan aptamer platform enabled unprecedented proteome breadth, it measures relative protein abundance and can include technical biases or off-target effects. Targeted validation of key findings by other proteomic platforms like immunoassays or mass spectrometry is essential. We also acknowledge that batch effects and demographic confounds, although mitigated by our harmonization strategy, may not be fully eliminated. Subtler sources of variability such as differences in comorbidities, or medications across sites could influence the plasma proteome. Additionally, certain groups, notably PDD and FTD, had smaller sample sizes, which limited power for discovery. Future consortia efforts should prioritize recruitment of these underrepresented subpopulations to fully characterize their proteomic profiles. Finally, our pathway and network analyses are inherently correlative. Follow-up mechanistic studies using human-specific *in vitro* models like stem cell-derived organoids are needed to determine causality and to explore whether modulating the identified pathways can modify disease outcomes.

Going forward, building on this work will entail not only validating the candidate biomarkers in independent cohorts but also leveraging our multi-omic, multi-cohort strategy to explore therapeutic interventions and extending the approach to pre-symptomatic or at-risk populations.

In summary, by integrating plasma proteomic data at an unprecedented scale across multiple diseases and sites, we have created a comprehensive map of systemic alterations in neurodegeneration. Our findings reveal a shared immune-metabolic dysfunction underlying AD and PD alongside distinct molecular signatures in disorders like ALS, and they highlight new biomarker candidates and druggable pathways for intervention. This work demonstrates the power of consortium-scale, harmonized proteomics to capture the multi-system nature of neurodegenerative diseases, bridging mechanistic insights to translational applications.

Ultimately, our integrative approach paves the way for improved blood-based biomarkers and therapeutic targets, moving the field closer to precision diagnosis and disease-modifying treatments for these devastating disorders.

## Resource Availability

### Materials availability

This study did not generate new materials.

## Data and code availability

The Global Neurodegeneration Proteomics Consortium dataset (v1) used and analysed in the current study is available through the Alzheimer’s Disease Data Initiative’s AD Workbench (https://discover.alzheimersdata.org/catalogue/datasets/e2f3536b-d97b-4303-89bd-6864200807a4)^17^. Researchers who wish to access this controlled dataset are required to register and submit a Data Use Agreement. Summary results generated in this manuscript are available in Supplementary Tables (S2-S6 and S14-S17).

## Acknowledgements

The authors are grateful to the cohort contributors, patients, donors, and families who helped to make this research possible. We are also thankful to Farhad Imam, Varsha Krish, and the team at Gates Ventures for their ongoing support with the Global Neurodegeneration Proteomics Consortium. This work was supported by the Australian Government’s Medical Research Future Fund MRF2040081 (C.A.F. and A.S.); philanthropic funding from Paul & Valeria Ainsworth family and Neil & Norma Hill Foundation (C.A.F.); the NIH R35NS132179 (J.D.R.), P30AG072973 (H.M.W., J.M.B., C.S., and R.H.S.), RF1AG064227 (C.S), R21TR003589 (H.M.W.), R01AG07816 (H.M.W.), U19AG068054 (H.M.W.), K23AG090757 (R.S.); Alzheimer’s Research UK (L.M.W.); Alzheimer’s Association 23AARG-1023 (H.M.W.), AARF-23-1145318 (RS); New Vision Research Charleston Conference on Alzheimer’s Disease (CCAD) 2024-001-1 (R.S.); American Academy of Neurology (R.S.), American Brain Foundation (R.S.), and Association for Frontotemporal Degeneration (R.S.); Wellcome Leap CARE (R.S.); Answer ALS Foundation (J.D.R.); and Robert Packard Center for ALS Research at Johns Hopkins University (J.D.R.). The funders of this work played no role in the design of the study, the running of experiments and analyses, the interpretation of the results, and the writing of the manuscript.

## Author contributions

A.S. conceived the paper and designed the study; A.S. performed the analyses on the GNPC cohort with input and feedback from L.A., L.M.W., J.V., M.W.L., and R.S.; C.A.F. and A.S. performed the enrichment analyses; H.M.W., J.M.B., R.H.S., and J.D.R. provided clinical neurology and diagnostic insights; C.A.F. and A.S. wrote the manuscript, which was edited and approved by all authors.

---

**Extended Data Figure 1.**
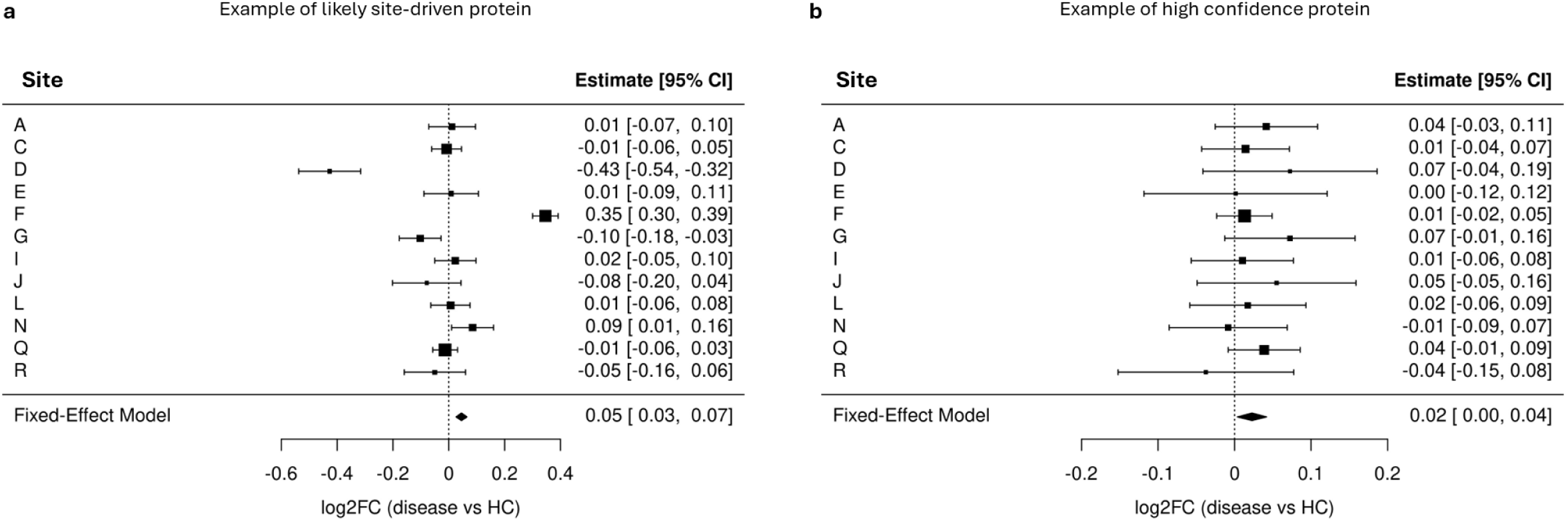
Examples of labelled proteins after quality control. (**a**) Example of a protein that was flagged as being likely site-driven. In this example, the protein’s effect size is being largely driven by Site F. (**b**) Example of a protein that was flagged as being high confidence. In this example, the direction and effect size are largely matched across the different contributing sites. Graphs show effect size (log2 fold change) and 95% confidence intervals. Abbreviations: CI: confidence interval; disease: Alzheimer’s disease; HC: healthy control.

**Extended Data Figure 2.**
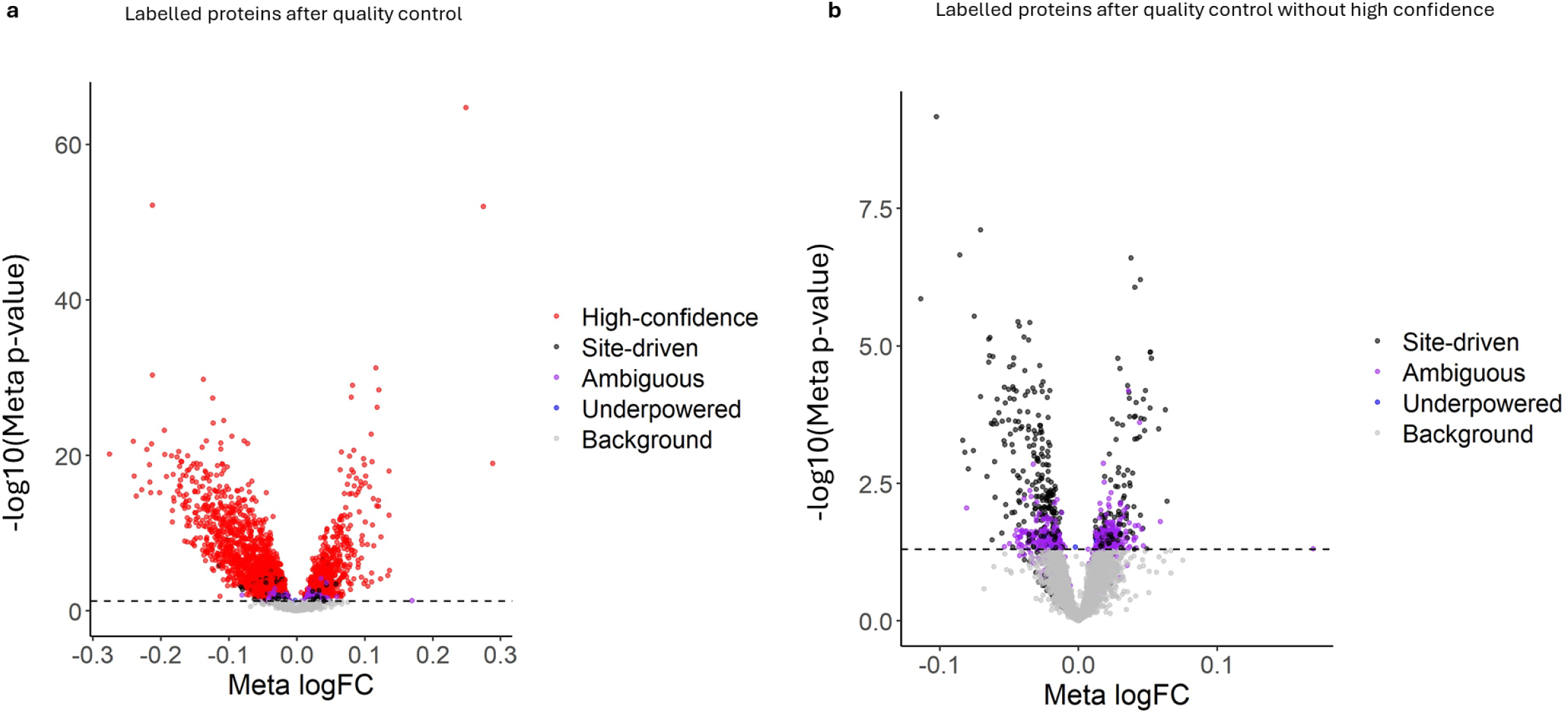
Alzheimer’s disease plasma proteins labelled following quality control checks. (**a**) Volcano plot shows labelled Alzheimer’s disease plasma proteins across high confidence, likely site-drive, ambiguous, underpowered, and background. (**b**) Volcano plot shows the same labelled Alzheimer’s disease plasma proteins with high confidence ones removed to improve the visualization of likely site-driven, ambiguous, underpowered, and background proteins. Both plots show meta p-values and fold change. Dotted black line represents *p* = 0.05.

